# Engineering phenotypic heterogeneity for functional organization in microbial populations

**DOI:** 10.64898/2026.09.14.751370

**Authors:** Marcela Villegas-Plazas, Irene Otero-Muras, Juan Nogales

## Abstract

Engineering microbial populations to perform complex functions requires programming not only cellular behaviour but also the functional organization through which biological activities are distributed across populations. While phenotypic heterogeneity is often regarded as variability to suppress, it can also serve as a foundation for organizing specialized functions within genetically homogeneous microbial systems. Here, we introduce PROMETEO, a modular genetic-circuit framework that programs population composition through architecture-encoded regulatory and translational asymmetries. Using a library of 27 asymmetric bistable circuits, we demonstrate that circuit architecture reproducibly specifies phenotypic distributions spanning a broad range of population compositions without continuous external induction. Programmed population structures remained stable over serial propagation and were qualitatively conserved across *Escherichia coli* and *Pseudomonas putida*. Stochastic and deterministic modelling accurately predicted architecture-dependent population compositions and hysteresis regimes, providing a quantitative framework for rational design. We further show that programmable population composition supports multiple modes of functional organization, including stable parallel specialization, inducible temporal redistribution of cellular states, and spatial ecological compartmentalization through biofilm-associated and planktonic subpopulations. As a demonstration of these capabilities, architecture- programmed organization enabled distributed Congo Red biotransformation through coordinated reductive and oxidative activities, achieving up to 98% dye removal in spatially compartmentalized populations. Together, these results establish population composition as a programmable property of genetic circuit design and provide a general strategy for engineering distributed functions within genetically homogeneous microbial populations.

## INTRODUCTION

Engineered microorganisms are emerging as programmable living technologies for sustainable manufacturing, environmental remediation, medicine, and biological information processing (*1*, *2*). As the functional scope of these applications expands, microbial systems are increasingly required to execute complex and often incompatible tasks, including multistep metabolic transformations, dynamic environmental adaptation, and coordinated regulatory responses. Advances in synthetic biology have enabled microorganisms to perform increasingly sophisticated biological functions through modular genetic design, mathematical modelling, and rational circuit engineering (*3*). As these functions become more numerous and interdependent, the central challenge is no longer only how to engineer functions, but how to engineer their functional organization across microbial populations in a predictable, stable, and controllable manner (*4*, *5*).

To address this challenge, two dominant engineering strategies have emerged. One relies on multitasking single-cell systems, in which multilayered genetic circuits dynamically coordinate multiple biological functions within individual cells (*6*, *7*). The other exploits synthetic microbial consortia, where complementary functions are distributed across specialized strains (*8*, *9*). Despite their different design strategies, both strategies ultimately aim to achieve functional organization at the population level by enabling distributed metabolism, coordinated regulation, and division of labor (*10*). Although these approaches have demonstrated considerable potential (*11*, *12*), each remains constrained by intrinsic limitations. Increasing circuit complexity imposes substantial metabolic burdens on multitasking single-cell factories (*13*, *14*), whereas synthetic consortia frequently suffer from unstable population dynamics and limited long-term controllability (*15*). These limitations have motivated growing interest in a third engineering strategy: achieving functional organization directly within genetically homogeneous populations, thereby combining the genetic simplicity of monocultures with the functional specialization typically associated with microbial communities.

Here, we define microbial functional organization as the controlled distribution of complementary biological functions across specialized functional subpopulations within a genetically homogeneous microbial population. Rather than representing undesirable biological noise, phenotypic heterogeneity is increasingly recognized as an engineering substrate that enables higher-order functional organization (*16–18*). Recent studies have shown that this potential can be harnessed through bistable phenotypic switching, population-partitioning systems (*10*) and, more recently, recombinase-based multistep cell-fate branching for precise control of population ratios (*19*). However, despite these advances, the ability to specify and stably maintain defined functional subpopulations within genetically homogeneous populations remains largely unexplored. Consequently, although phenotypic heterogeneity can now be engineered, harnessing programmable population composition as a predictable and stable design principle for engineering functional organization remains an outstanding challenge.

Among the regulatory architectures developed in synthetic biology, toggle switches (TS) have emerged as the canonical framework for establishing bistable cell states (*20*) and are increasingly recognized as fundamental regulatory modules underlying cellular decision-making in gene regulatory networks (*21*). By establishing two mutually exclusive expression states, these circuits provide the regulatory basis for cellular differentiation, memory formation, and stable phenotypic commitment (*22*). Their bistable dynamics are characterized by hysteresis, whereby the switching threshold depends on the previous state of the system (*23*). Narrow hysteresis regimes facilitate reversible transitions, whereas broader hysteresis landscapes promote persistent memory and stronger phenotypic commitment (*24*). Beyond their well-established applications in cellular decision-making, differentiation, and memory (*25–28*), TS are particularly attractive for engineering functional organization because they establish stable functional subpopulations capable of supporting complementary biological functions within isogenic populations (*10*, *29–31*). However, exploiting these dynamical properties to engineer functional organization in a predictable and controllable manner remains a major challenge.

Classical TS designs consist of two mutually repressing transcriptional regulators that generate two alternative stable equilibrium states (*26*). State transitions are typically induced by chemical inducers that transiently relieve repression and drive the system across a bifurcation threshold (*27*). Switching may be reversible or irreversible, thereby determining the persistence of cellular memory following inducer removal (*32*). While these designs provide robust control over individual cell states, they rely on graded inducer inputs to influence state occupancy, resulting in poorly predictable population compositions and limiting their ability to engineer functional organization.

Collectively, these studies establish bistable regulatory circuits as tunable systems in which circuit design governs state stability and switching behavior (*23*, *27*, *33*, *34*). However, most existing TS architectures continue to rely on inducer titration to influence the proportion of cells occupying each phenotypic state, making population composition highly sensitive to environmental fluctuations and difficult to maintain over time. Consequently, their practical applications have largely focused on temporal programming of cellular states (*35–37*), rather than on stable parallel functional organization within microbial populations. This limitation highlights a critical gap in the current synthetic biology toolbox: the lack of autonomous strategies that establish population composition as a programmable design variable for engineering functional organization.

To address this challenge, we introduce PROMETEO (PROgrammable MEtabolic heTErOgeneity), a design framework that engineers functional organization by programming population composition in isogenic microbial populations. Unlike classical inducible bistable systems, PROMETEO specifies population composition through circuit design rather than inducer titration. Rather than determining population composition through inducer dosage, PROMETEO encodes it directly into circuit architecture through regulatory and translational asymmetries while retaining inducible state switching when external control is desired. As a result, circuit design becomes the primary determinant of population composition, creating a design space that couples programmed population composition with hysteresis-driven state redistribution. To facilitate implementation, PROMETEO is organized as a modular Golden Gate Level 1 library for rapid circuit assembly and deployment.

Here, we combine systematic experimental characterization with mathematical modelling to map the design space of PROMETEO and establish programmable population composition as a design principle for engineering functional organization. We identify regulatory asymmetry as the design principle through which circuit design specifies population composition. We demonstrate three distinct modes of functional organization enabled by programmable population composition: temporal switching between functional states, stable parallel functional partitioning within isogenic populations, and spatial lifestyle differentiation into planktonic and biofilm-associated subpopulations. Finally, we demonstrate this design principle in *Pseudomonas putida* KT2440 by coordinating the distributed biotransformation of Congo Red through anaerobic reductive and aerobic oxidative reactions that are physiologically incompatible within individual cells.

## RESULTS

### Construction of a systematic design space of asymmetric toggle switches

To test whether regulatory asymmetry can intrinsically program population composition, we developed PROMETEO, a framework based on engineered TS architectures designed to determine how regulatory asymmetry programs population composition within isogenic bacterial populations. Our aim was to embed programmable population composition directly into circuit architecture, thereby specifying population composition as an intrinsic property of circuit design without reliance on continuous external induction. Each PROMETEO architecture builds on the canonical double-negative feedback motif defining TS topology, in which two transcriptional repressors mutually inhibit each other. For broad applicability and to minimize regulatory cross-talk, we selected three repressor systems, BetI, TtgR, and TetR, each responsive to a distinct small-molecule inducer: choline (cho), naringenin (nar), and anhydrotetracycline (aTc), respectively. These components, derived from the engineered Marionette platform (*38*), provide well-characterized regulatory modules with distinct basal leakage levels, dynamic ranges and inducibility profiles, parameters known to strongly influence bistable dynamics and topology-dependent state behaviour (*20*, *34*, *39*).

We combined orthogonal repressor systems to generate a systematic landscape of mutually repressive bistable configurations. While all circuits share the same TS topology, they differ in their regulatory configuration, defined by the specific pairing of repressors and their target promoters. These architectures were organized into three core configurations: configuration 1, BetI–TtgR; configuration 2, BetI–TetR; and configuration 3, TtgR–TetR (Fig. 1A). To introduce controlled regulatory asymmetry, we systematically varied ribosome binding site (RBS) strengths at each regulatory node using three well-established parts: SBGst (weak, W), B0034 (medium, M), and BCD8 (strong, S) (Fig. 1B). This combinatorial design yielded 27 bistable configurations spanning a defined design space encompassing a broad range of regulatory and translational imbalances. Each construct is therefore defined by a unique combination of repressor identity and translational strength, together specifying its programmed population composition.

**Figure 1.**
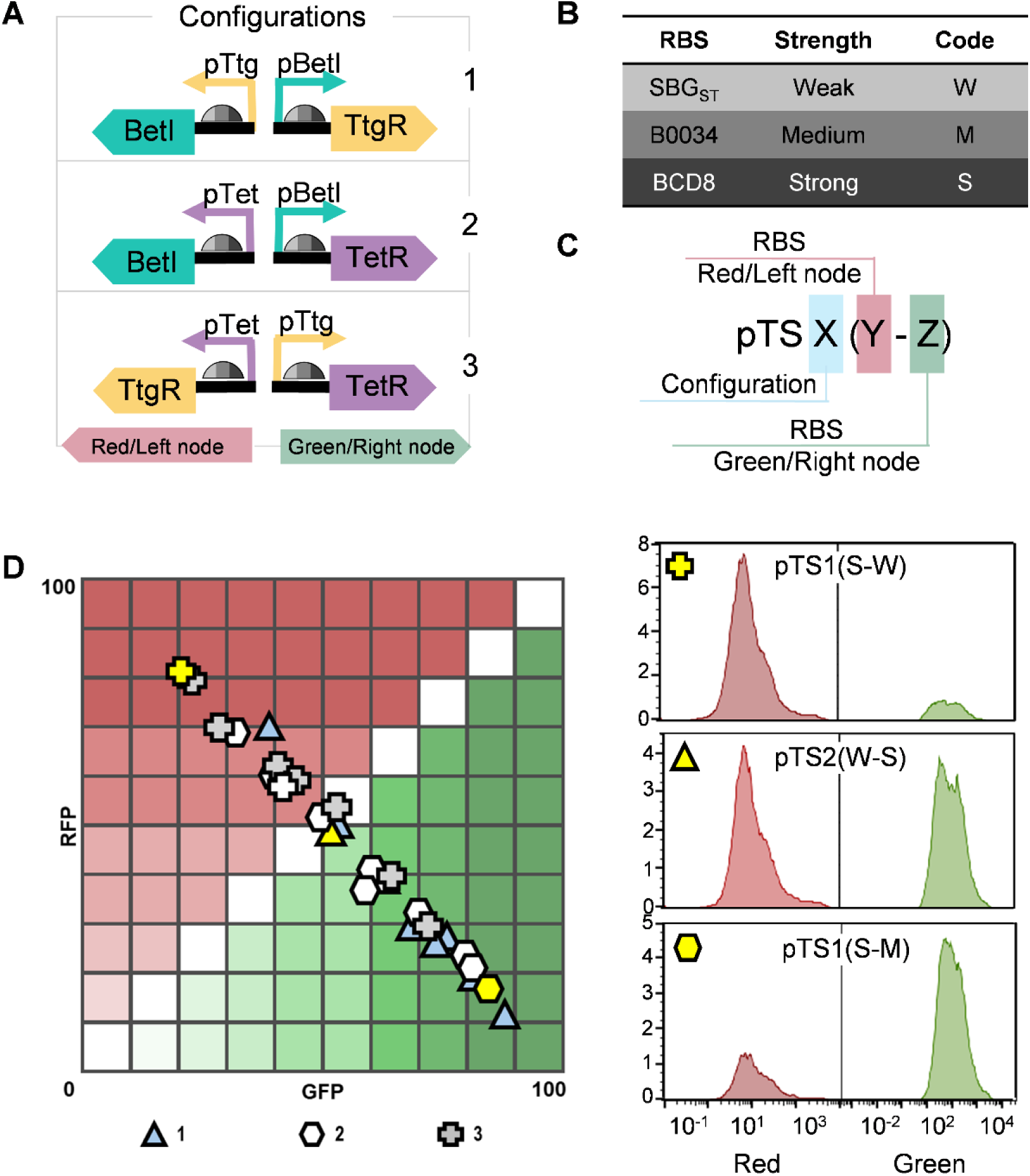
Architectural design and population composition landscape of the PROMETEO library. (A) Three bistable toggle-switch (TS) configurations comprising different pairs of mutually inhibitory transcriptional regulators: Configuration 1 (BetI– TtgR), Configuration 2 (BetI–TetR) and Configuration 3 (TtgR–TetR). The left node controls RFP expression and the right node GFP expression. (B) Ribosome binding sites (RBSs) used to generate weak (W), medium (M) and strong (S) translational outputs. (C) PROMETEO nomenclature. Variants are designated as pTSX(Y–Z), where X indicates the circuit configuration and Y and Z the RBS strengths of the left/red and right/green nodes, respectively. (D) Population composition landscape of the PROMETEO library under non-induced conditions. Grid coordinates indicate the relative abundance of Red- and Green-state subpopulations determined by flow cytometry. Representative single-cell fluorescence distributions for selected variants are shown on the right. Histogram y-axis values indicate cell counts (×10²).

For standardized referencing, each construct was named in the format pTSX(Y-Z), where X denotes the configuration (1–3), and Y and Z indicate the RBS strength (W, M, S) at the red/left and green/right nodes, respectively (Fig. 1C). For example, pTS2(M-S) designates a circuit from configuration 2 with a medium RBS on the left node and a strong RBS on the right. RBS strength categories are defined by calibrated translational output in *Pseudomonas putida KT2440*, our primary deployment chassis in this study. When reporting *Escherichia coli* data, we retain the same labels to preserve one-to-one traceability across hosts. All constructs were assembled using MoClo-compatible standards (Supplementary Table S1). Together, this library establishes an experimental design space for investigating how circuit architecture programs population composition.

### Circuit architecture programs population composition in *E. coli*

To test whether circuit architecture systematically programs population composition, we evaluated how different circuit configurations influence the resulting population structure in *E. coli,* a host in which the regulatory components and translational parts used in PROMETEO are well characterized (*38*). To do so, we constructed reporter versions of each circuit, placing mRFP1 and GFPmut3 on the left and right nodes, respectively. This configuration enabled single-cell resolution of mutually exclusive expression states, with red- and green-fluorescent subpopulations reporting activation of each regulatory node. We then screened the full set of 27 bistable configurations in *E. coli* DH5α using flow cytometry, quantifying red/green population distributions to determine whether circuit- encoded regulatory asymmetry could systematically program population composition (Fig. 1D).

*E. coli* overnight cultures were diluted into fresh medium and sampled during exponential phase, yielding >50,000 single-cell measurements per sample. Across the library, all architectures exhibited robust bimodal distributions, confirming that bistability was preserved across diverse translational configurations (Fig. 1D). Notably, the relative occupancy of red- and green-expressing states remained highly reproducible across biological replicates (Supplementary Table S2), demonstrating that circuit architecture reproducibly programs population composition.

The PROMETEO library programmed a broad spectrum of population compositions, with individual configurations tiling the state space from near-complete Red-state occupancy to near-complete Green-state occupancy (Fig. 1D). These results demonstrate that circuit design is sufficient to reproducibly program these distributions across a broad design space, without reliance on chemical induction. Collectively, these findings establish circuit architecture as a sufficient design principle for programming population composition in bistable circuits.

### Architecture-programmed population composition is conserved across host contexts

To determine whether architecture-programmed population composition is conserved across physiological contexts, we transferred PROMETEO to *P. putida* KT2440, a metabolically versatile chassis widely used in synthetic biology and industrial biotechnology. This transition enabled evaluation of whether bistability, population composition, and long-term state occupancy remained robust under a distinct physiological background.

All 27 circuits maintained reproducible bimodal population distributions in this host, consistent with partitioning into two phenotypic subpopulations. To evaluate the long-term stability of the resulting population compositions, each construct was propagated through at least six serial transfers, with Red/Green ratios quantified at stationary phase in every passage (Fig. 2A, Supplementary Table 3). The resulting population compositions remained highly reproducible across all configurations, with less than 4– 8% variation between passages, demonstrating that the encoded population compositions remained stable across serial propagation despite physiological and nutritional variation. These results show that encoded population structures can persist over extended propagation, indicating that programmed population composition can be specified as a stable property of circuit design. To interpret host-dependent effects on circuit behaviour, we quantified promoter strength, relative RBS performance and basal leakage for the BetI-, TtgR- and TetR-based regulatory systems in *P. putida* KT2440 (Fig. 2B). These measurements provided a chassis-specific framework for interpreting architecture-dependent population behaviours across hosts.

**Figure 2.**
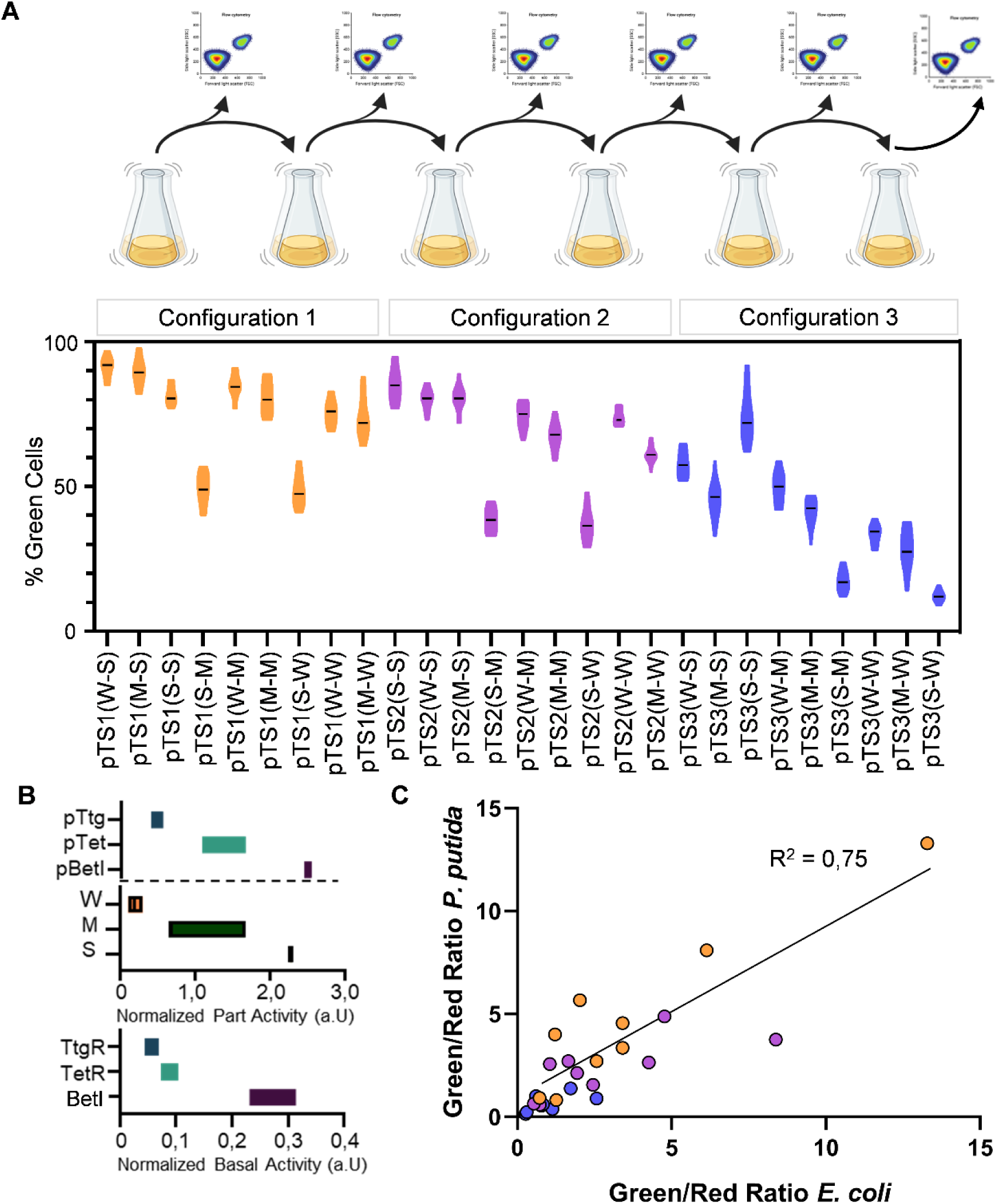
Stable and host-conserved population composition across the PROMETEO library. (A) Stability of programmed population compositions across the PROMETEO library during six successive stationary-phase cultures in *P. putida* KT2440 under non-induced conditions. Violin plots show the percentage of Green-state cells for each architecture from three independent biological replicates. (B) Quantitative characterization of the regulatory components in *P. putida* KT2440. Top panels show normalized promoter activities and RBS strengths. Bottom panel shows normalized basal leakage of the BetI-, TetR- and TtgR-based regulatory systems. (C) Comparison of Green/Red population ratios measured for each PROMETEO architecture in *E. coli* DH5α and *P. putida* KT2440. Each point represents one architecture; the solid line indicates linear regression (R² = 0.75).

Direct comparison of *E. coli* and *P. putida* KT2440 revealed a strong overall correlation in variant behaviour across hosts (R² = 0.75), despite moderate shifts in absolute population composition (Fig. 2C). Circuits generating near-balanced populations in *E. coli* remained similarly balanced in *P. putida*, whereas architectures biased towards one phenotypic state preserved the same directional preference. Thus, host context primarily rescales population proportions without altering the qualitative relationship between circuit architecture and programmed population composition. The largest deviations were associated with constructs carrying weaker RBSs, consistent with the known host dependence of translational parts in the Golden Standard system (*40*). Together, these results demonstrate that the relationship between circuit architecture and population- level organization is qualitatively conserved across distinct bacterial hosts, despite substantial differences in host physiology.

### Mathematical modelling predicts architecture-programmed population composition

Having established that circuit architecture reproducibly programs population composition, we next sought to understand how architecture-encoded regulatory and translational asymmetries quantitatively determine population composition. We therefore developed a stochastic model based on partial integro-differential equations (PIDE) that describes the temporal evolution of phenotypic state distributions across the population. Rather than focusing on individual cellular trajectories, the framework captures how the distribution of phenotypic states evolves over time, ultimately shaping the emergence and stabilization of population-level organization. Model outputs were directly mapped onto GFP and RFP fluorescence distributions measured by flow cytometry, enabling quantitative comparison between predicted and experimentally observed population distributions. A detailed description of the modelling framework, governing equations and simulation parameters is provided in Supplementary Note 1 and Supplementary Tables S4 and S5.

The model accurately recapitulated the experimentally observed population distributions under non-induced conditions (Fig. 3A). When the strongest RBS (BCD8) was placed at the GFP/right node and the weakest site (SBGst) at the RFP/left node, the fraction of Green-state cells at stationary phase increased to 91%, 83% and 60% for configurations 1, 2 and 3, respectively (Fig. 3A). Reversing the translational asymmetry redistributed the population towards the Red state, reducing Green-state fractions to 55%, 30% and 12%, in close agreement with the experimentally observed fluorescence distributions (Fig. 2A).

**Figure 3.**
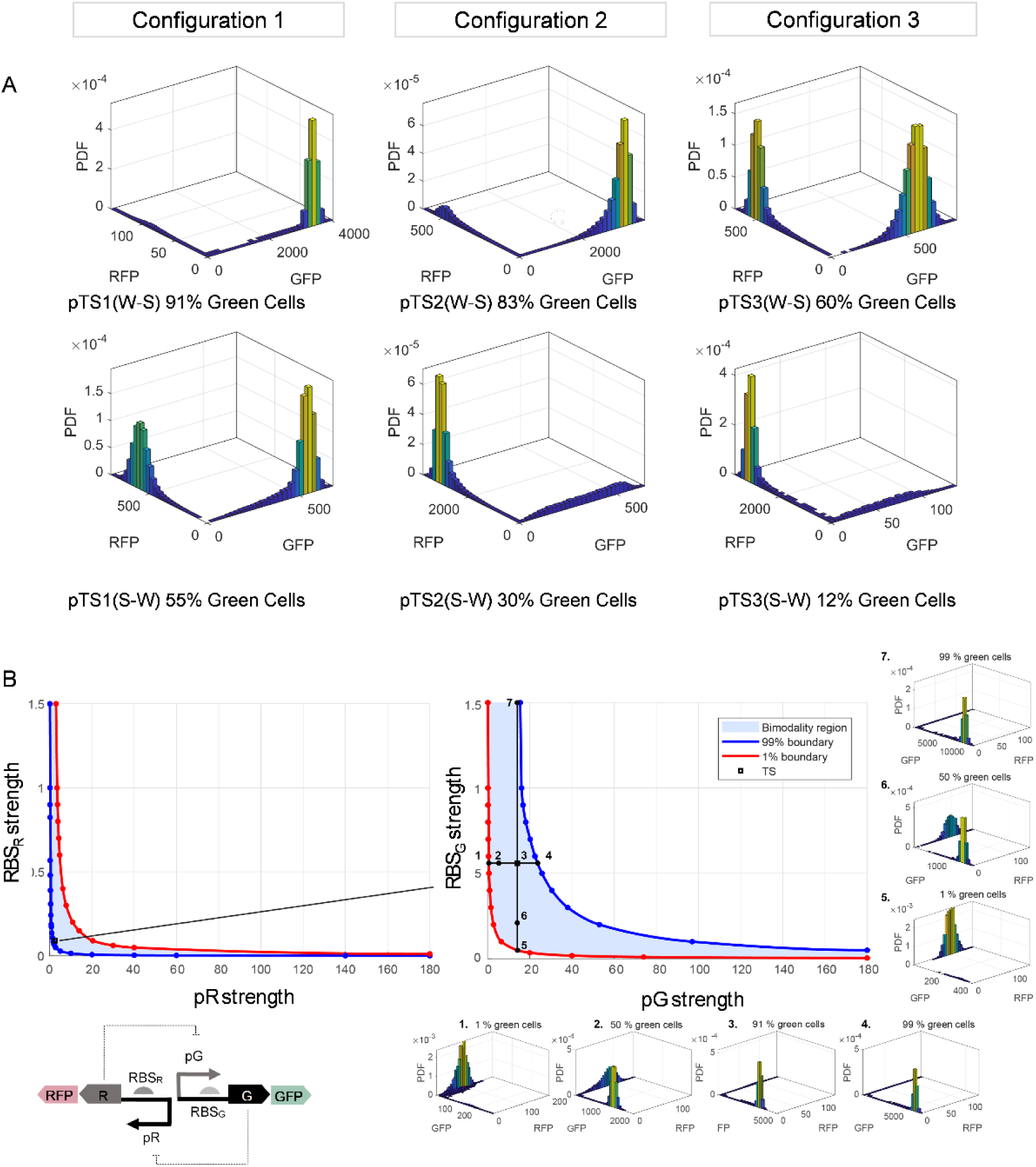
Population-level modelling predicts architecture-programmed population composition. (A) Simulated probability density distributions of GFP and RFP expression across the three PROMETEO configurations under opposing translational asymmetry regimes. Top panels correspond to architectures carrying weak Red-node and strong Green-node RBS combinations (W–S), whereas bottom panels correspond to the inverse asymmetry (S–W). Percentages indicate the simulated fraction of Green-state cells. (B) Parameter-space analysis of bistable population organization. Blue-shaded regions indicate parameter combinations supporting bimodal population distributions. Blue and red boundary curves correspond to 99% and 1% Green-state occupancy, respectively. Left panel, effects of Red-node promoter (pR) and RBS strength. Right panel, effects of Green-node promoter (pG) and RBS strength. Insets show representative simulated population distributions spanning the full range of Green- state occupancy.

We next used the model to predict how circuit parameters reshape programmable population composition by systematically mapping bimodal probability distributions across the parameter space defined by promoter and RBS strengths (Fig. 3B). This analysis identified the asymmetry regimes supporting stable population compositions.

The resulting landscape revealed that population composition is both tunable and quantitatively predictable across defined asymmetry regimes. For a fixed circuit configuration, modulation of either promoter or RBS strength alone was sufficient to redistribute population composition from near-complete Red-state occupancy to near-complete Green-state occupancy. For example, systematic variation of promoter strength (states 1 → 2 → 3 → 4, Fig. 3B) or RBS strength (states 5 → 6 → 3 → 7, Fig. 3B) progressively shifted state occupancy across the full dynamic range. Importantly, the landscape provides practical design rules for tuning population composition through promoter or RBS engineering, while also identifying circuit architectures that expand the bistable regime and support broader ranges of stable population structures. Together, these results establish a predictive framework through which circuit architecture can be used to rationally program population composition.

### Circuit architecture shapes hysteresis-driven state switching and memory

Having established that circuit architecture predictably programs population composition, we next investigated whether it also controls the dynamics of transitions between alternative population states through hysteresis. Configurations with narrow hysteresis ranges favour reversible switching and rapid redistribution between phenotypic states, whereas broader hysteresis ranges reduce reversibility, thereby reinforcing state commitment and promoting more persistent phenotypic memory (Fig. 4A).

**Figure 4.**
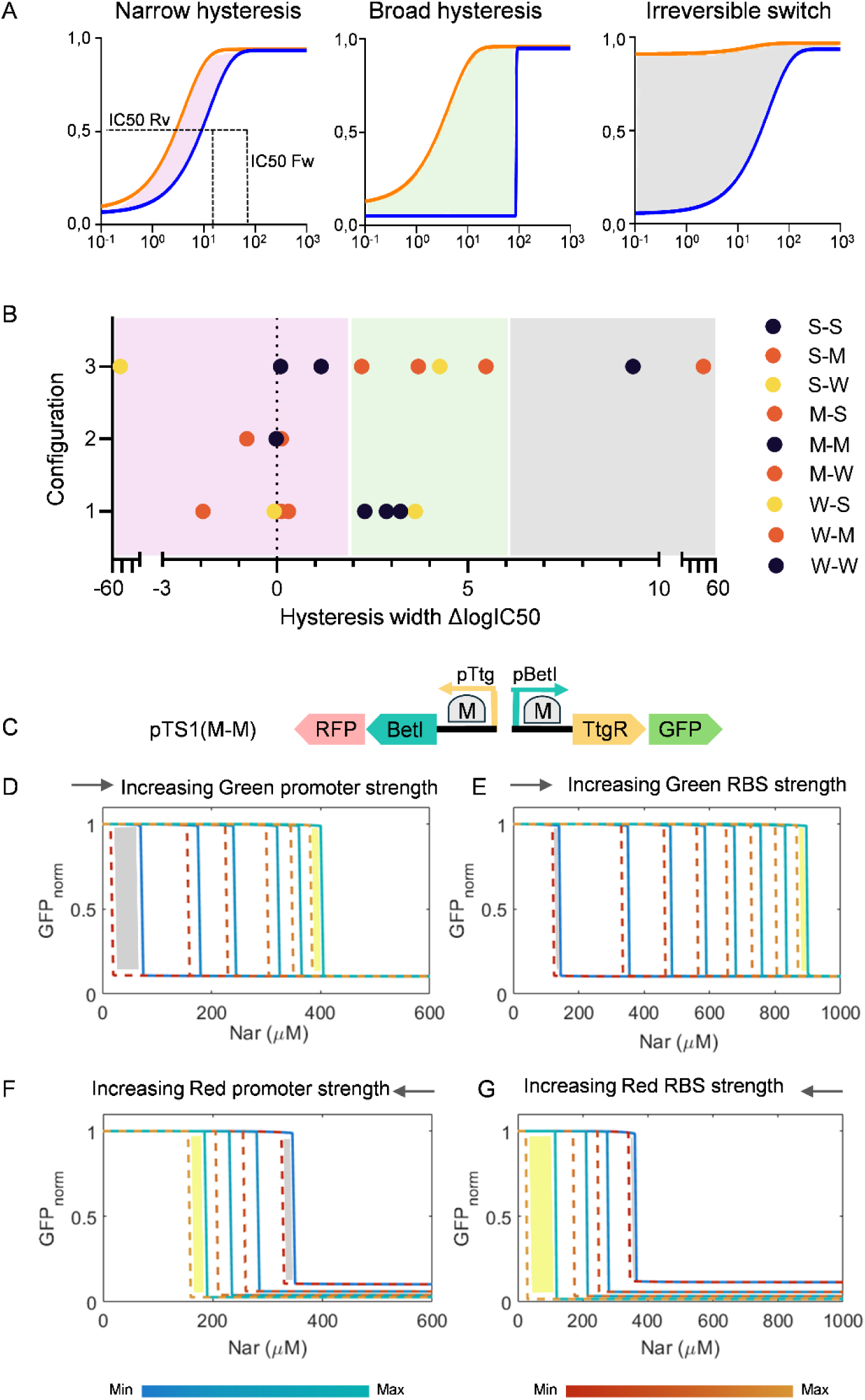
Regulatory asymmetry shapes hysteresis landscapes across the PROMETEO framework. (A) Representative narrow, broad and effectively irreversible hysteresis regimes. Hysteresis width was quantified as ΔlogIC50 between forward (blue) and reverse (orange) switching thresholds. (B) Distribution of hysteresis widths (ΔlogIC50) across PROMETEO variants. Points are coloured according to the Red/Green RBS combination. (C) Representative toggle-switch architecture used for modelling analyses (pTS1(M–M)). (D–G) ODE-derived hysteresis landscapes showing the effects of increasing Green-node promoter strength (D), Green-node RBS strength (E), Red-node promoter strength (F) and Red-node RBS strength (G) on switching thresholds. Solid and dashed curves represent forward and reverse transitions, respectively.

To quantitatively compare memory behaviour across the PROMETEO library, we characterized hysteresis in all 27 variants in *P. putida* KT2440 (Supplementary Fig. 1). Forward and backward induction curves were obtained for each construct, and hysteresis width was quantified as the difference in log10 midpoints (ΔlogIC50) (Fig. 4B, Supplementary Table S6). Variants with ΔlogIC50 ≈ 0 displayed narrow, near-reversible hysteresis regimes and therefore supported rapid transitions between alternative phenotypic states. Intermediate values (ΔlogIC50 ≈ 2–6) generated extended bistable regimes in which switched populations remained committed at inducer concentrations substantially lower than those required for activation, consistent with strong phenotypic memory. At the extreme, variants with ΔlogIC50 ≥ 9, or lacking detectable reversible transitions, approached effectively irreversible switching behaviour under the tested conditions, consistent with highly persistent phenotypic memory.

Distinct hysteresis regimes emerged across the three regulatory configurations. Configuration 1 (BetI–TtgR) generated heterogeneous switching behaviours spanning both reversible and extended-memory regimes. Representative variants such as pTS1(W–S), pTS1(S–S), pTS1(W–W) and pTS1(M–M) displayed broad hysteresis (ΔlogIC50 > 2; Fig. 4B), consistent with strong persistence of the induced state. In contrast, variants including pTS1(M–S) and pTS1(W–M) remained close to reversible switching regimes, exhibiting ΔlogIC50 values near 0 and therefore rapid redistribution between alternative phenotypic states.

Configuration 2 (BetI–TetR) constrained nearly all variants to narrow hysteresis ranges (ΔlogIC50 ≈ 0), resulting in predominantly reversible switches. This behaviour was particularly evident in pTS2(M–S), in which hysteresis collapsed towards symmetric switching, consistent with rapid responsiveness while retaining minimal detectable memory (Fig. 4B).

In contrast, Configuration 3 generated the broadest hysteresis regimes and strongest memory effects across the PROMETEO framework. Multiple asymmetric RBS combinations, including representative variants such as pTS3(W–S), pTS3(M–S), pTS3(W–M) and pTS3(S–M), occupied extended bistable regimes (ΔlogIC50 > 3; Fig. 4B), consistent with highly persistent phenotypic commitment. At the extreme, effectively irreversible switching regimes emerged. Notably, pTS3(M–W) displayed ΔlogIC50 values above 45, whereas pTS3(S–W) exhibited no detectable reverse transition, indicating loss of accessible state reversion under the tested conditions (Fig. 4B). These observations indicate that regulatory asymmetry influences not only population composition but also the accessibility of transitions between alternative states.

To mechanistically interpret the architecture-dependent hysteresis and memory regimes observed experimentally, we developed an ODE-based mean-field model that captures representative switching and hysteretic dose–response behaviours across the three configurations. A detailed description of the model and parameters is provided in Supplementary Note 2 and Supplementary Table S7.

Using this framework, we simulated the hysteresis behaviour of the representative variant pTS1(M–M) while systematically varying promoter and translational parameters. The model quantitatively recapitulated the major hysteresis regimes observed experimentally across asymmetry regimes. Increasing promoter or RBS strength on the Green-node shifted switching thresholds towards higher inducer concentrations (Fig. 4D-E), consistent with the broad hysteresis observed in asymmetric variants such as pTS3(W–M) → pTS3(W–S) or pTS3(M-M) → pTS3(M-S) (Supplementary Fig. 1C). Conversely, increasing promoter or RBS strength on the opposing Red-node shifted switching thresholds towards lower inducer concentrations while broadening the hysteresis loop (Fig. 4F-G), as observed in the sequence pTS3(W-W) → pTS3(M–W) → pTS3(S–W) (Supplementary Fig. 1C), progressively shifting the system towards effectively irreversible state commitment (Fig. 4G). In the model, this transition occurs when the lower bifurcation point of the bistable regime moves into the negative orthant, corresponding to non-physical inducer concentrations and thereby eliminating accessible reverse transitions.

Together, these results demonstrate that regulatory asymmetry defines a continuum of hysteresis and memory regimes, ranging from highly reversible state redistribution to extended bistable commitment and effectively irreversible transitions. Thus, beyond programming population composition, circuit architecture establishes a second programmable layer of control by governing the dynamical accessibility of transitions between alternative phenotypic states, from reversible redistribution to effectively irreversible commitment.

### Programmable population organization enables multiple strategies for distributed metabolic function

To determine whether architecture-programmed population organization can coordinate complex metabolic functions within an isogenic population, we applied PROMETEO to Congo Red (CR) degradation, a model process requiring sequential reductive and oxidative reactions with distinct physiological requirements (*41*). These coupled transformations are physiologically difficult to sustain efficiently within individual cells, making CR degradation dependent on coordinated metabolic specialization (*42*, *43*). As a result, degradation is typically achieved more efficiently by microbial consortia than by monocultures, for which coordinating incompatible metabolic states remains challenging despite extensive metabolic engineering efforts (*44*, *45*). CR degradation therefore provides a stringent functional test for whether programmable population organization can coordinate environmentally distinct metabolic functions within a single genetic background.

The sequential nature of this pathway provides an opportunity to evaluate not only whether distributed metabolism can be achieved, but also how different organizational strategies influence its execution. Stable population composition enables persistent allocation of complementary functions across differentiated subpopulations, whereas inducible state redistribution enables dynamic rebalancing of those functions over time. Together, these complementary organizational modes demonstrate how circuit architecture can implement multiple strategies for coordinating distributed metabolism within genetically homogeneous populations.

### Intrinsic population composition enables distributed metabolic coordination

CR degradation proceeds through two main steps: reductive cleavage of the azo bond by AzoR, followed by oxidative detoxification of the resulting aromatic amines by the CotA laccase. We therefore constructed TS_CR_(I) by placing *azoR* and *cotA* on opposite arms of pTS3(M–S) (Supplementary Table S8, Fig. 5A), a circuit that generates an approximately balanced population composition (∼50:50). In this design, one subpopulation expresses *AzoR* whereas the complementary subpopulation expresses *CotA*, thereby embedding complementary metabolic functions directly into circuit architecture without requiring external induction.

**Figure 5.**
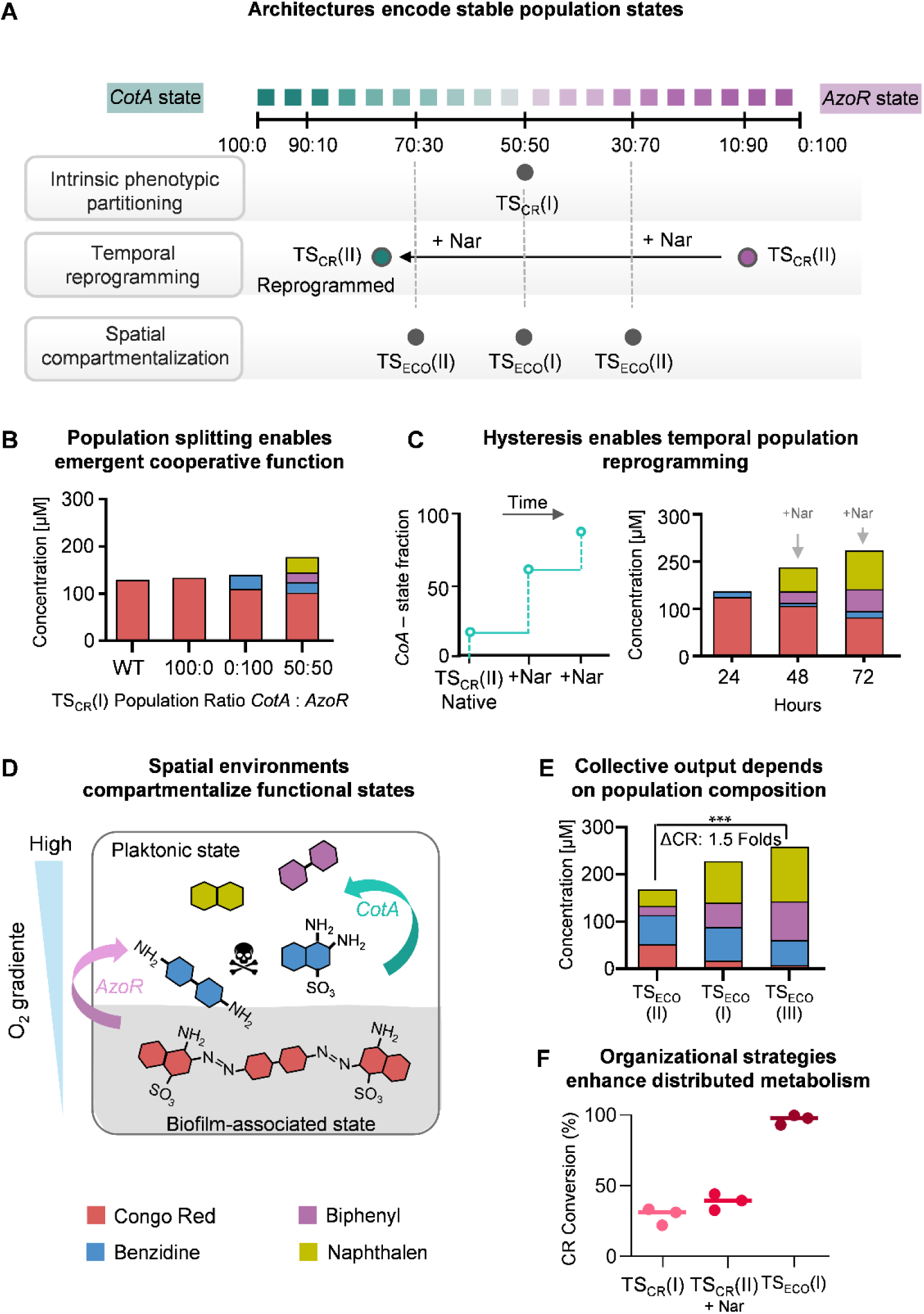
Programmable population organization enables multiple strategies for distributed metabolism. (A) PROMETEO-derived application constructs used for distributed metabolism. TS_CR_(I) and TS_CR_(II) implement intrinsic population composition and inducible temporal redistribution, respectively, whereas TS_ECO_ variants couple architecture-programmed population composition to spatial ecological compartmentalization. (B) HPLC analysis of Congo Red (CR) transformation after 72 h by TS_CR_(I) populations with different CotA:AzoR population compositions. (C) Temporal redistribution of TS_CR_(II) populations following sequential Nar induction and corresponding metabolite profiles after 24, 48 and 72 h. (D) Schematic of the TS_ECO_ strategy. AzoR is coupled to the biofilm-associated state through co-expression with *YedQ*, whereas CotA is associated with the planktonic state. (E) Metabolite profiles of TS_ECO_ variants carrying different intrinsic CotA:AzoR population ratios. (F) Comparison of CR conversion achieved using intrinsic population composition TS_CR_(I), inducible temporal redistribution TS_CR_(II) and spatial ecological compartmentalization TS_ECO_(I). Points represent independent biological replicates; horizontal bars indicate the mean.

We benchmarked this organizational strategy against the corresponding single-function states. Cultures of *P. putida* harbouring TS_CR_(I) were compared under three conditions: (i) uninduced bifurcated populations (∼50:50 *AzoR*:*CotA*), (ii) a *CotA*-only state induced with aTc (0.00025 mM) and (iii) an AzoR-only state induced with Nar (1 mM). All cultures were incubated with 140 µM CR and monitored by HPLC for 72 h.

Neither single-enzyme condition achieved substantial CR removal (<10%), whereas bifurcated TS_CR_(I) populations removed up to ∼30% of CR (Fig. 5B). Notably, oxidative products such as naphthalene and biphenyl emerged only after 48 h (Supplementary Fig. S2), indicating that oxidative detoxification followed prior AzoR-mediated azo bond reduction performed by the complementary subpopulation. This delayed appearance of oxidative products revealed a metabolic handoff emerging from intrinsic population composition. Thus, intrinsic population composition was sufficient to coordinate sequential metabolic functions without external induction.

### Inducible temporal reprogramming orchestrates sequential metabolic coordination

Although intrinsic population composition in TSCR(I) successfully coupled the reductive and oxidative branches of the pathway, the delayed appearance of oxidative products (Supplementary Fig. S2) suggested that pathway performance remained limited by the temporal coordination of these complementary metabolic activities. We therefore designed TS_CR_(II) to dynamically reprogram population composition over the course of the degradation process (Fig. 5A), allowing the dominant metabolic activity to shift as pathway requirements changed. TS_CR_(II) was based on pTS1(W–S), a circuit exhibiting a strong intrinsic asymmetry (∼90% AzoR, ∼10% CotA; Supplementary Table S8) that initially biases the population towards the reductive state and promotes early azo cleavage. Prior characterization of the hysteresis landscape of pTS1(W–S) (Supplementary Fig. 1A, Fig. 5C) enabled rational selection of inducer regimes that progressively shifted the population towards the oxidative state. Accordingly, cultures received a partial Nar induction after 24 h in CR-containing medium, followed by a second, stronger induction at 48 h to complete the transition towards *CotA*-dominated populations. This strategy enabled controlled navigation through the bistable landscape, progressively reallocating metabolic activities in accordance with the sequential demands of the pathway.

This inducible temporal reprogramming outperformed static organizational strategies. After 72 h, TS_CR_(II) populations achieved ∼40% CR degradation, exceeding both single-enzyme states (<10%) and intrinsically bifurcated TS_CR_(I) populations (∼30%) (Fig. 5C). Moreover, benzidine accumulation, which persisted under static conditions, was nearly eliminated. HPLC analysis revealed more efficient progression through the degradation pathway, with increased accumulation of downstream oxidative products such as biphenyl and naphthalene. Together, these results indicate that temporal reprogramming of population composition improves the coordination of sequential metabolic functions by aligning population organization with the changing functional demands of the pathway. In this configuration, circuit architecture was used not only to establish functional specialization but also to dynamically reorganize metabolic activities as the pathway progressed.

### Programmable population organization drives lifestyle differentiation and spatial structuring

Although temporal redistribution improved pathway coordination, both TS_CR_(I) and TS_CR_(II) operate within physically homogeneous conditions. This represents a potential limitation for CR degradation, as the two complementary activities are favoured by distinct physiological environments. *AzoR*-mediated azo reduction is promoted under oxygen-limited conditions, whereas *CotA*-dependent oxidation requires molecular oxygen for activity (*46*, *47*). We therefore reasoned that further improvements in pathway coordination might require not only functional specialization, but also the capacity to organize distinct cellular states into physically differentiated environments.

To determine whether circuit architecture could translate programmed population composition into lifestyle differentiation and spatial structuring, we developed TS_FILM_ (TS- driven Framework for Induced Lifestyle Modulation), a functional extension of PROMETEO in which activation of the Green state drives co-expression of *GFPmut3* and *YedQ*, a diguanylate cyclase that elevates intracellular c-di-GMP and promotes biofilm formation (*48*). The opposing Red state, reported by RFP and lacking *YedQ* expression, represents the alternative non-biofilm-forming population. Three TS_FILM_ variants were constructed from pTS3(M–S), pTS3(M–W), and pTS2(W–W), corresponding to TS_FILM_(I), TS_FILM_(II), and TS_FILM_(III), respectively (Fig. 6A). These architectures encode approximately 50:50, 70:30, and 30:70 Red:Green population compositions (Supplementary Table S8), enabling evaluation of how programmed population composition influences lifestyle allocation and subsequent spatial structuring.

**Figure 6.**
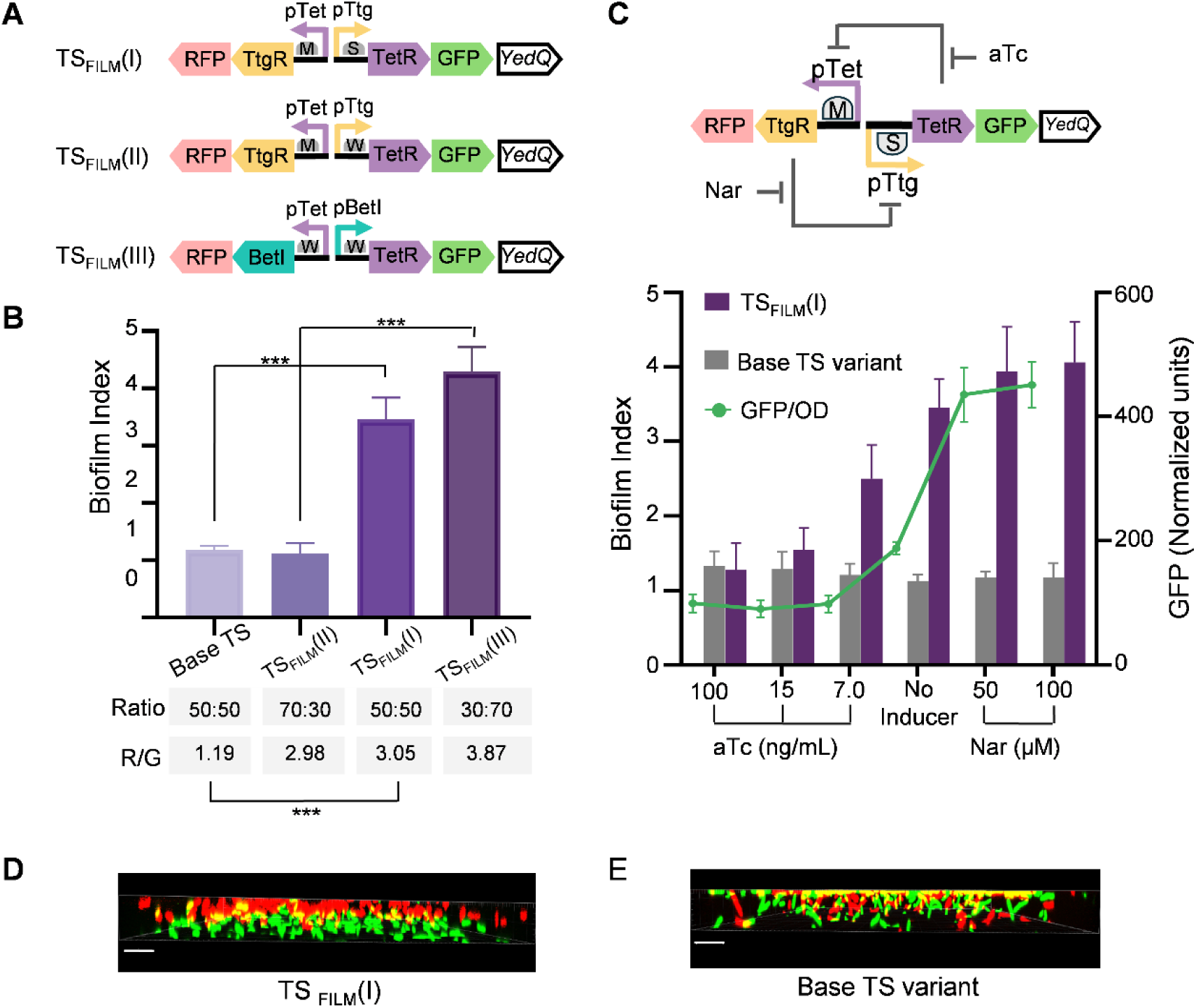
Toggle-controlled lifestyle differentiation generates programmable spatial organization. (A) Genetic architectures of the TS_FILM_ constructs, in which the diguanylate cyclase *YedQ* was coupled to the GFP-associated toggle state to promote c-di-GMP-dependent biofilm formation. Three variants encode distinct intrinsic Red:Green population compositions (70:30, 50:50 and 30:70). (B) Biofilm formation after 48 h of static incubation for the TS_FILM_ variants and the corresponding Red/Green fluorescence ratios measured in the planktonic fraction. Bars represent mean ± s.d. (n = 3 biological replicates). ***P < 0.001 (one-way ANOVA with Tukey’s multiple-comparison test). (C) Inducible modulation of TS_FILM_(I) with Nar or aTc. Biofilm index (bars) and normalized GFP fluorescence (green line) are shown for each induction condition. Bars represent mean ± s.d. (n = 3 biological replicates). (D) Confocal image of TS_FILM_(I) after static growth showing vertical stratification of Green and Red populations. (E) Confocal image of the corresponding TS control lacking *YedQ*. Yellow regions indicate overlap between red and green fluorescence. Scale bars, 20 μm.

To determine how programmed population composition translated into lifestyle allocation, TS_FILM_ variants were assayed after 48 h of static incubation. Biofilm formation, measured by crystal violet staining, scaled with the fraction of Green-state cells encoded by each architecture, demonstrating that population-level behaviour can be quantitatively programmed through circuit architecture (Fig. 6B). Normalization to total cell density confirmed that the observed differences reflected programmed population composition rather than changes in overall biomass accumulation. Consistent with their programmed population compositions, TS_FILM_(I) (≈50:50) produced intermediate biofilm levels, TS_FILM_(II) (≈70:30 Red:Green) generated reduced biofilm formation approaching the baseline TS control lacking *YedQ*, whereas TS_FILM_(III) (≈30:70 Red:Green) yielded the strongest biofilm signal, consistent with enrichment of the adherent subpopulation. Flow cytometry of the planktonic fraction revealed RFP/GFP distributions consistent with preferential depletion of Green-state cells from the suspension and their corresponding enrichment within the biofilm-associated fraction (Fig. 6B).

We next evaluated inducibility using TS_FILM_(I). Addition of Nar, which progressively shifts the toggle towards the Green node, increased both GFP output and biofilm formation in a dose-dependent manner, thereby increasing the abundance of the biofilm-forming subpopulation (Fig. 6C). In contrast, the control TS lacking *YedQ* displayed increased GFP levels without a corresponding increase in biofilm formation. Thus, biofilm modulation resulted from coupling programmed population composition to *YedQ* expression rather than from toggle induction alone. Conversely, addition of aTc, which biases the system towards the Red node, reduced GFP levels in both the control and TS_FILM_(I). In TS_FILM_(I), this shift was accompanied by reduced biofilm formation to baseline levels comparable to the control circuit lacking *YedQ* (Fig. 6C).

To directly visualize spatial organization, we performed confocal microscopy on static cultures (Fig. 6D–E). TS_FILM_(I) populations formed a vertically stratified structure in which Green-state, YedQ-expressing cells formed a biofilm layer attached to the bottom surface of the well, whereas Red-state cells remained predominantly in the overlying planktonic phase (Fig. 6D, Supplementary Video 1). In contrast, the baseline TS lacking *YedQ* produced a diffuse red–green mixture with no detectable segregation (Fig. 6E, Supplementary Video 2). Together, these observations demonstrate that programmed population composition can be translated into inducible lifestyle differentiation and vertical spatial structuring within an isogenic microbial population, thereby establishing physically distinct cellular niches that provide the structural basis for subsequent ecological specialization.

### Programmable spatial organization enables ecological metabolic compartmentalization

Having established that programmable spatial organization generates physically distinct cellular niches, we next asked whether these structures could be exploited to achieve ecological compartmentalization. Although temporal reprogramming improved pathway performance, it remained constrained by the distinct physiological requirements of the two complementary reactions: AzoR-mediated azo reduction is favoured under oxygen-limited conditions, whereas CotA-dependent oxidation requires molecular oxygen for activity. We therefore hypothesized that spatially organized biofilm and planktonic compartments would provide distinct physiological environments capable of supporting the simultaneous execution of both reactions within an isogenic population.

To implement this concept, we engineered TS_ECO_ (TS-driven <u>E</u>cological <u>Co</u>mpartmentalization), a spatially organized system derived from the TS_FILM_ framework. *AzoR* was co-expressed with the diguanylate cyclase *YedQ* on the right node, coupling reductive metabolism to the biofilm-associated compartment, whereas CotA was expressed in the opposing planktonic compartment (Fig. 5D). TS_ECO_ variants were generated from previously characterized architectures exhibiting intrinsic CotA:AzoR bifurcation ratios of approximately 50:50, 70:30, and 30:70, corresponding to TS_ECO_(I), TS_ECO_ (II), and TS_ECO_ (III), respectively (Fig. 5A). By varying the programmed population composition, this series tested how different architectural configurations influence compartmentalized CR transformation.

Performance analysis revealed a strong dependence on the underlying programmed population composition. Progressive enrichment of the AzoR-associated biofilm compartment led to markedly enhanced degradation efficiency. In particular, TS_ECO_(III), enriched in the AzoR-expressing biofilm compartment (∼30:70 CotA:AzoR), achieved up to 98% CR removal after 72 h (Fig. 5E), substantially exceeding both the intrinsically bifurcated TS_CR_(I) (∼30%) and the temporally reconfigured TS_CR_(II) system (∼40%) (Fig. 5F). Although absolute benzidine levels appeared comparable across TS_ECO_ variants, normalization to total CR degradation revealed a substantial reduction in intermediate accumulation in TS_ECO_(III), consistent with more efficient downstream oxidation into biphenyl and naphthalene. These results indicate that ecological metabolic compartmentalization improves pathway flux by reducing intermediate accumulation while enabling complementary metabolic activities to proceed simultaneously under their preferred physiological conditions.

Importantly, these degradation levels substantially exceeded those of previously reported single-strain systems under comparable conditions and approached the degradation efficiencies typically associated with engineered microbial consortia (*41*, *49–51*). Collectively, these findings demonstrate that programmable spatial organization enables ecological metabolic compartmentalization within genetically homogeneous populations, thereby allowing ecologically specialized subpopulations to coordinate distributed metabolism.

## DISCUSSION

### Circuit architecture as a programmable determinant of population organization

Synthetic biology increasingly seeks to distribute biological functions across specialized cellular states, particularly through microbial consortia and multicellular regulatory strategies (*20*, *52*). However, achieving stable and tunable coordination within genetically homogeneous populations remains considerably less developed. Toggle switches have classically been used to control transitions between individual cellular states, supporting memory storage and inducible regulation, but their potential to specify how those states are distributed across a population has remained comparatively underexplored (*26*, *27*). Here, we show that principles of population organization can themselves be encoded as a circuit-level design property through systematic modulation of regulatory and translational asymmetries in bistable architectures.

Across the PROMETEO library, differences in promoter identity, regulatory leakage and translational strength generated reproducible and stable population compositions spanning a broad continuum of phenotypic ratios (Fig. 1D, Fig. 2A). These distributions emerged under non-induced conditions and remained stable over serial propagation, indicating that population composition can be specified intrinsically by circuit architecture rather than maintained through continuous environmental control. Thus, bistable circuits can function not only as inducible memory devices, but also as architectures that stabilize defined phenotypic state distributions across genetically homogeneous populations.

The conservation of architecture-to-phenotype relationships across *E. coli* and *P. putida* further suggests that programmable population organization is governed by regulatory design principles that extend beyond a single host context (Fig. 2C). Although absolute population ratios differed quantitatively between chassis, the directional biases imposed by specific regulatory configurations were strongly preserved. Together with the PIDE- based modelling framework (Fig. 3), these observations support a predictable relationship between engineered circuit asymmetry and population composition. More broadly, they indicate that population organization can arise as an intrinsic and qualitatively conserved consequence of regulatory architecture, rather than solely as a response to external environmental inputs.

Recent advances have demonstrated precise control of cell-type ratios through recombinase-mediated differentiation and lineage branching architectures (*19*). In contrast, PROMETEO does not prescribe discrete lineage outcomes, but programs the occupancy, stability, and accessibility of alternative phenotypic states within a common genetic background. This distinction shifts the engineering objective from specifying the abundance of predefined cell types to engineering the organizational principles through which functional specialization emerges and reorganizes within isogenic populations.

### Circuit architecture couples population composition and organizational dynamics

Programming population organization requires control not only over how phenotypic states are distributed, but also over how stably those distributions are maintained and how readily they can be reorganized. In PROMETEO, population composition, state persistence and inducible redistribution were jointly modulated within the same circuit-design space. Variations in promoter and translational strength reshaped both the relative occupancy of toggle states and the stability of transitions between them (Fig. 3B, Fig. 4B). Thus, bistability operates here not merely as a mechanism for binary switching, but as a design property that couples population composition with the dynamics through which organizational states are maintained or reconfigured (*20*).

The hysteresis landscapes of the PROMETEO library defined a continuum of organizational regimes, from readily reversible state redistribution to strongly persistent or effectively irreversible population configurations (Fig. 4B). Narrow hysteresis supported greater accessibility between phenotypic states, whereas broader hysteresis stabilized committed states and constrained their subsequent redistribution (*53*). The ODE-based model linked promoter and translational parameters to bistable landscapes and switching thresholds at the mean-field level (Fig. 4D–G), while the PIDE framework connected these same design regimes to population-level state occupancy and distributional dynamics (Fig. 3, Supplementary Note 1). Together, these models indicate that circuit architecture specifies not only which population compositions are favoured, but also the dynamical accessibility of transitions between them.

This coupling between composition and dynamics enabled distinct modes of functional coordination. In TS_CR_(I), architecture-defined population composition maintained stable specialization between reductive and oxidative subpopulations, giving rise to a metabolic relay without externally imposed redistribution (Fig. 5B). By contrast, TS_CR_(II) exploited the hysteretic regime of pTS1(W–S) to reassign population states through staged inducer inputs, converting intrinsic specialization into an externally orchestrated sequence of metabolic functions (Fig. 5C). These two regimes illustrate that circuit architecture can be selected either to preserve functional organization or to permit its controlled temporal reconfiguration. Population composition and organizational dynamics are therefore not independent circuit outputs, but coupled and jointly engineerable properties of bistable design.

### Programmable population organization extends across spatial and ecological dimensions

Once population composition and organizational dynamics can be programmed, differentiated states can be coupled to distinct physiological roles and arranged across higher levels of functional organization. PROMETEO therefore extends beyond controlling phenotypic ratios or temporal state redistribution: by linking circuit-defined states to alternative microbial lifestyles, it enables functional coordination across compositional, temporal, spatial and ecological dimensions. In this framework, population organization does not merely distribute cellular states, but determines how complementary functions are assigned, positioned, and coordinated within a genetically homogeneous population.

The TS_FILM_ and TS_ECO_ systems illustrate how this principle can be extended from state allocation to spatially structured ecological specialization. Coupling toggle states to biofilm and planktonic lifestyles generated spatially segregated subpopulations occupying distinct physicochemical environments. TS_ECO_ subsequently exploited this architecture-programmed spatial organization by associating *AzoR* expression with the biofilm-enriched compartment and *CotA* expression with the planktonic compartment, thereby positioning reductive and oxidative activities within physiologically differentiated niches (Fig. 5D). Biofilm-associated cells provided hypoxic conditions favourable for azo reduction, whereas planktonic cells supported oxidative detoxification in the surrounding oxygenated environment. Spatial organization thus became a functional component of pathway design rather than a passive consequence of growth.

Viewed together, the PROMETEO configurations represent distinct organizational strategies rather than independent circuit applications. Intrinsic population composition supported stable functional specialization, inducible state redistribution enabled temporal coordination, and lifestyle differentiation introduced spatial and ecological compartmentalization. These regimes demonstrate that complementary metabolic functions can be coordinated through multiple dimensions of population organization while remaining encoded within a common circuit architecture. Importantly, this coordination was achieved without assembling multispecies consortia or relying on dedicated intercellular signalling systems.

Whereas recombinase-based differentiation frameworks generate predefined cellular lineages with user-defined abundance (*19*), PROMETEO exploits architecture- programmed bistable state organization to couple population composition with temporal, spatial, and ecological modes of coordination.

The enhanced Congo Red degradation achieved by TS_ECO_(III), reaching approximately 98% removal after 72 h (Fig. 5E), provides a functional demonstration of this organizational logic. The significance of this result lies less in the degradation pathway itself than in showing that architecture-programmed spatial differentiation can generate cooperative metabolic behaviour typically associated with microbial consortia or naturally structured communities (*50*). Ecological specialization therefore emerges here not as a passive response to environmental heterogeneity, but as an engineered consequence of population organization.

### Toward population organization as a programmable design variable

The findings presented here support a broader engineering role for bistable genetic circuits than has traditionally been recognized. Rather than serving primarily as devices for controlling binary cellular decisions and inducible memory, toggle architectures can be engineered to specify how functional organization emerges across genetically homogeneous populations. PROMETEO illustrates this transition by showing that population composition, organizational dynamics and ecological specialization arise as coupled consequences of circuit architecture, collectively redefining the design space accessible through bistable genetic systems.

Viewing circuit architecture as a determinant of population organization expands the engineering possibilities available for microbial systems. Population composition, temporal reorganization and ecological compartmentalization become engineerable organizational properties rather than isolated circuit behaviours, enabling distributed functions to be coordinated across multiple levels of biological organization. In this framework, phenotypic heterogeneity is no longer viewed primarily as stochastic variability to suppress, but as the biological substrate through which programmable population organization can be engineered. This perspective shifts the engineering objective from specifying the behaviour of individual cells toward specifying the functional organization of entire microbial populations.

Beyond its engineering implications, this framework also provides an experimentally tractable platform for investigating how collective organization emerges from coordinated phenotypic differentiation within genetically homogeneous systems. Although these findings should not be interpreted as establishing a new form of multicellularity, they demonstrate that many organizational properties typically associated with microbial consortia or spatially structured communities can instead be rationally encoded within a single isogenic population through circuit architecture. More broadly, these principles position population organization as a programmable engineering design variable that complements traditional approaches focused on controlling individual cellular behaviour, thereby expanding the conceptual and practical design space for synthetic microbial systems.

Despite these promising advances, translating programmable population organization beyond laboratory settings will require understanding how architecture-defined population compositions behave under prolonged cultivation, variable environmental conditions, and application-specific functional burdens. Future efforts should also address the long-term evolutionary stability of programmed population structures and explore how population-level organization can be integrated with sensing and feedback-control systems to enable adaptive redistribution of specialized functions. Nevertheless, the results presented here establish that population composition can be encoded as an intrinsic property of circuit architecture, providing a general framework for engineering distributed functions within genetically homogeneous microbial populations and extending microbial design beyond the optimization of population-average behaviours.

## MATERIALS AND METHODS

### Bacterial strains and plasmid construction

All cloning procedures were performed in *Escherichia coli* DH5α (New England BioLabs). Constructs were subsequently transferred into *Pseudomonas putida* KT2440 for experimental implementation due to its metabolic robustness, biofilm-forming capacity, and compatibility with synthetic circuits. Strains were cultured in LB medium supplemented with appropriate antibiotics: ampicillin (Amp, 100 µg mL⁻¹), kanamycin (Km, 50 µg mL⁻¹), or gentamycin (Gm, 10 µg mL⁻¹). All toggle switch assays were conducted in M9 minimal medium (1× M9 salts, 2 mM MgSO₄, 0.1% (w/v) casamino acids, 0.0005% (w/v) thiamine) with 0.2% (w/v) glucose as the carbon source. To enhance growth without obscuring bistable behaviour, media were supplemented with 10% (v/v) LB.

Plasmids were assembled using the Golden Standard modular cloning system (*40*). Level 0 parts, including ribosome binding sites (RBSs), terminators, fluorescent proteins, and linkers, were obtained from the SBG collection. Promoters and regulators (BetIR, TtgR, and TetR) were amplified from the Marionette sensor collection (*38*) using oligonucleotides listed in Supplementary Table S1. Restriction sites BsaI and BpiI were domesticated as needed. The *YedQ* diguanylate cyclase was PCR-amplified from pSYedQ plasmid (*48*), and *AzoR* from *P. putida* KT2440 genomic DNA, while *CotA* was codon-optimized for *Pseudomonas*, domesticated, and synthesized by GenScript Biotech Corporation.

### Toggle Switch design and library assembly

Each variant in the PROMETEO toggle switch (TS) library was constructed using a modular architecture composed of four genetic modules: two mutually inhibitory regulatory modules and two fluorescent sensor modules. Regulatory modules contained repressor genes (tetR, betI or ttgR) expressed from promoters repressed by the opposing regulator, thereby forming a double-negative feedback loop. Distinct ribosome binding sites (RBSs) were incorporated to generate translational asymmetry across circuit variants.

Sensor modules encoding GFPmut3 and mRFP1 were placed under the control of opposing repressor-responsive promoters to report alternative circuit states. These reporters were included only in characterization constructs.

All four modules were assembled into Level 2 plasmids using the Golden Standard system (*40*) and cloned into RK2-based broad-host-range vectors. For application-specific constructs, TS architectures lacking fluorescent reporters were reformatted into Level 1 plasmids using customized fusion linkers flanked by Golden Standard fusion sites 4–5. Functional modules encoding *YedQ*, *AzoR* and *CotA* were subsequently incorporated to generate TS_FILM_, TSCR and TS_ECO_ application constructs.

All genetic modules and assembled plasmids are listed in Supplementary Table S1.

### Phenotypic characterization via flow cytometry and fluorometry

#### Flow cytometry analysis of intrinsic phenotypic partitioning

To assess architecture-encoded phenotypic distributions in *E. coli* and *P. putida*, single colonies of each TS variant were cultured overnight in supplemented M9 medium. Cultures were diluted to an OD₆₀₀ of 0.1 and grown to stationary phase at 37 °C (*E. coli*) or 30 °C (*P. putida*). Cells were collected, washed, and diluted in PBS to OD₆₀₀ = 0.2. Flow cytometry was performed using a MACSQuant™ VYB cytometer (Miltenyi Biotec), acquiring 50,000 events per sample. GFP was excited at 488 nm and detected using a 525/40 nm filter, whereas mRFP was excited at 561 nm and detected using a 615/20 nm filter. Single-cell gating was performed using FSC/SSC parameters. Fluorescence gates were defined using single-color positive controls generated from the corresponding promoter–fluorophore combinations of each regulatory system and were maintained constant across experiments and biological replicates. Data were analyzed using FlowJo v10 and are reported as the mean of three biological replicates.

#### Fluorometric Hysteresis Assays

To characterize hysteretic behaviour, TS strains were cultured in 96-well plates containing 300 µL per well of M9 medium supplemented with concentration gradients of Nar or aTc. Initial Red- and Green-state populations were established by overnight preconditioning with saturating concentrations of aTc (100 ng mL⁻¹) or Nar (1 mM), respectively. Cells were subsequently washed twice with inducer-free M9 medium prior to transfer into induction gradients. Cultures were incubated at 30 °C in a VICTOR Nivo™ plate reader (PerkinElmer), and OD₆₀₀, GFP, and RFP fluorescence were recorded automatically every 30 min over 24 h. GFP fluorescence was measured using 480/30 nm excitation and 530/30 nm emission filters, whereas RFP fluorescence was measured using 580/20 nm excitation and 625/30 nm emission filters. Plates were subjected to orbital shaking at 600 rpm for 15 s before and after each measurement, as well as at 15- min intervals between measurements. Dose–response curves were generated from endpoint fluorescence measurements and used to calculate switching thresholds and hysteresis widths (ΔlogIC50).

### Biofilm quantification and spatial imaging

Biofilm formation was quantified using a crystal violet (CV) staining assay adapted from Benedetti et al. (2016). Cultures were grown statically in 96-well plates containing 300 µL of M9 medium supplemented with inducers where indicated. For induction assays, aTc was used at 7, 15 and 100 ng mL⁻¹, and Nar at 50 and 100 µM. OD₆₀₀ and fluorescence were monitored every 30 min for 48 h using a VICTOR Nivo™ plate reader, without shaking. After incubation, supernatants were collected for flow cytometric analysis of planktonic fractions. Wells were subsequently washed and stained with 200 µL of 0.1% (w/v) CV for 30 min, followed by three washes and solubilization with 150 µL of 33% (v/v) acetic acid. Absorbance at 590 nm was recorded, and the biofilm index was calculated as CV absorbance normalized to total biomass (OD₆₀₀).

For confocal imaging, cultures were grown statically for 24 h in µ-Slide Well ibiTreat chambers under the same culture conditions used for biofilm assays. Immediately before imaging, cultures were immobilized by adding low-melting-point agarose to a final concentration of 0.4% (w/v). Z-stack fluorescence imaging was performed using a Leica TCS SP5 multispectral confocal system. GFP and RFP were excited at 488 nm and 561 nm, respectively. Image stacks were processed using Imaris Viewer, and vertical fluorescence distributions were analyzed to assess spatial segregation between biofilm-associated and planktonic subpopulations.

### Metabolic coordination assays and metabolite profiling

Distributed metabolic coordination was evaluated using Congo Red (CR) as a model sequential redox substrate. Cultures were grown statically at 30 °C in 20 mL M9 medium supplemented with 0.2% glucose and 140 µM CR, with initial inoculation at OD₆₀₀ = 0.2. For temporal redistribution experiments, cultures were induced with Nar at 24 h and 48 h to final concentrations of 340 µM and 600 µM, respectively. Samples were collected every 24 h, frozen until analysis, mixed 1:1 with methanol, and filtered through hydrophobic PTFE syringe filters (0.22 µm pore size, 13 mm diameter; SFPT-122) before HPLC analysis.

Metabolite profiling was performed using an Agilent 1260 Infinity II HPLC system equipped with an InfinityLab Poroshell 120 EC-C18 column (4.6 × 150 mm) and precolumn (4.6 × 5 mm). Samples were injected at 20 µL and separated at 1.0 mL min⁻¹ using water with 0.1% formic acid as solvent A and methanol with 0.1% formic acid as solvent B. The gradient was run from 80:20 A:B to 5:95 A:B, followed by re-equilibration to 80:20 A:B. CR, benzidine, biphenyl and naphthalene were quantified from peak areas using external calibration curves generated from analytical standards. Compounds were identified by retention time and quantified using diode array detection across 218–338 nm, encompassing the absorbance maxima of the analyzed metabolites.

### Modeling of phenotypic distribution

The stochastic population model was simulated using the IDESS Toolbox (*54*) which includes GPU-parallelized implementations of the Stochastic Simulation Algorithm (SSA) (*55*) and the PIDE semi-Lagrangian simulation solver implemented in SELANSI (*56*). Simulations of the deterministic ODE model were performed using MATLAB’s ode45 solver. Parameter identification was performed by maximum likelihood estimation using the enhanced Scatter Search (eSS) algorithm (*57*) or hybrid optimization. The observation of hysteresis indicates persistent bistable behaviour at the population level despite intrinsic noise. In the thermodynamic limit, stochastic fluctuations become less influential relative to system size, and the system approaches a mean-field description of bistable dynamics. Consistently throughout this work, “bistable” refers both to dose-response behaviour with two stable steady states and to bimodal population distributions corresponding to two underlying attractors.

All computations were performed in MATLAB R2024a on a Windows 11 workstation equipped with a 13th Gen Intel® Core™ i7-13700H processor and an NVIDIA GeForce RTX 4060 GPU with 3,072 CUDA cores.

Custom MATLAB scripts used in this study are openly available at: https://github.com/SBGlab/PROMETEO

### Statistical analysis

Unless otherwise stated, all experiments were performed using at least three independent biological replicates. Data are presented as mean ± standard deviation (SD). Statistical significance was assessed using one-way or two-way ANOVA followed by Tukey’s multiple-comparison test, as appropriate. Statistical analyses were performed using GraphPad Prism 9, with significance thresholds set at p < 0.05.

## FUNDING

This work was supported by the European Union’s Horizon 2020 Research and Innovation Programme under Grant Agreement No. 101027389 (MENTHOL) and by the Horizon Europe programme under Grant Agreement No. 101081782 (deCYPher). Additional support was provided by the Spanish Ministry of Science, Innovation and Universities (AEI/10.13039/501100011033) through grant PID2022-139247OB-I00 (Rob3D), and by the Spanish National Research Council (CSIC) through project LINCG25043 (FLAVO-GREEN). IOM acknowledges funding from the CellWise (ERC- 2024-COG-101170783) Contract of the European Union and AIA2025-164235-C44 project funded by MICIU/AEI/10.13039/501100011033/

## AUTHOR CONTRIBUTIONS STATEMENT

M.V.P. and J.N. conceived the study. M.V.P. performed the experiments. I.O.M. performed the mathematical modeling. M.V.P., I.O.M., and J.N. analyzed the data. M.V.P. and J.N. wrote the original draft. M.V.P., I.O.M., and J.N. reviewed and edited the manuscript.

## COMPETING INTERESTS

The authors declare no competing interests.

## DATA, CODE, AND MATERIALS AVAILABILITY

Custom MATLAB scripts and modeling resources used in this study are publicly available on GitHub: https://github.com/SBGlab/PROMETEO. Raw experimental data supporting all figures and supplementary figures, including flow cytometry datasets, HPLC chromatograms, plate-reader measurements, confocal microscopy data, plasmid sequences, and source data tables, have been deposited in Zenodo and are available at DOI: https://doi.org/10.5281/zenodo.20815606

Raw experimental data supporting all figures and supplementary figures, including flow cytometry datasets, HPLC chromatograms, plate-reader measurements, confocal microscopy data, and source data tables, have been deposited in Zenodo and are available at DOI: https://doi.org/10.5281/zenodo.20815606

## Notes

https://doi.org/10.5281/zenodo.20815606

### Competing Interest Statement

The authors have declared no competing interest.

